# Diagnostic and Prognostic Implications of Ribosomal Protein Transcript Coordination in Human Cancers

**DOI:** 10.1101/167940

**Authors:** James M. Dolezal, Arie P. Dash, Edward V. Prochownik

## Abstract

Ribosomes, the organelles responsible for the translation of mRNA, are comprised of rRNA and ~80 ribosomal proteins (RPs). Although canonically assumed to be maintained in equivalent proportions, some RPs have been shown to possess differential expression across tissue types. Dysregulation of RP expression occurs in a variety of human diseases, notably in many cancers, and altered expression of some RPs correlates with different tumor phenotypes and patient survival. To investigate the impact of global RP transcript (RPT) expression patterns on tumor phenotypes, we analyzed RPT expression of ~10,000 human tumors and 700 normal tissues with *t*-distributed stochastic neighbor embedding (*t*-SNE). We show here that normal tissues and cancers possess readily discernible RPT expression patterns. In tumors, this patterning is distinct from normal tissues, distinguishes tumor subtypes from one another, and in many cases correlates with molecular, pathological, and clinical features, including survival. Collectively, RPT expression can be used as a powerful and novel method of tumor classification, offering a potential clinical tool for prognosis and therapeutic stratification.

## Introduction

Eukaryotic ribosomes are among the most highly evolutionarily conserved organelles, comprised of four ribosomal RNAs (rRNAs) and approximately 80 ribosomal proteins (RPs). Responsible for translating mRNA into proteins, ribosomes were long believed to be nonspecific “molecular machines” with unvarying structures and function in different biological contexts. Recent evidence has shown, however, that some RPs are expressed in tissue-specific patterns and can differentially contribute to ribosome composition, affect rRNA processing, and regulate translation (Xue and Barna 2012). Despite the complexity of RP assembly in ribosomes, early studies of ribosome function revealed that the catalytic activity responsible for peptide bond formation might depend only on the presence of rRNAs and a small number of core RPs (Noller et al. 1992). This finding, in conjunction with the observation that some RPs are expressed in a tissue-specific manner, has led some to speculate that one purpose for the evolutionary emergence of RPs may have been to confer translational specificity and adaptability to ribosomes (Xue and Barna 2012; Guimaraes and Zavolan 2016).

An increasing body of evidence continues to show that RPs do, in fact, have an important role in imbuing ribosomes with mRNA translation specificity. During embryonic development, RPs are expressed at different levels across tissue types, and loss of RPs due to mutation or targeted knockdown produces specific developmental abnormalities in plants, invertebrates, and vertebrates. The tissue-specific patterning that occurs as a consequence of individual RP loss suggests that some RPs serve to guide the translation of specific subsets of transcripts in order to influence cellular development. Although the mechanism(s) by which RPs confer translation specificity are not entirely known, one may involve the alteration of ribosome affinity for transcripts with specific *cis*-regulatory elements, including internal ribosome entry sites (IRES) elements and upstream open reading frames (uORFs). (Xue and Barna 2012)

RPs also participate in a variety of extra-ribosomal functions. In normal contexts, ribosome assembly from rRNAs and RPs is a tightly regulated process, with unassembled RPs undergoing rapid degradation. Disruption of ribosomal biogenesis by any number of extracellular or intracellular stimuli induces ribosomal stress, leading to an accumulation of unincorporated RPs. These free RPs are then capable of participating in a variety of extra-ribosomal functions, including the regulation of cell cycle progression, immune signaling, and cellular development. Many free RPs bind to and inhibit MDM2, a potentially oncogenic E3 ubiquitin ligase that interacts with p53 and promotes its degradation. The resulting stabilization of p53 triggers cellular senescence or apoptosis in response to the inciting ribosomal stress. Additional extra-ribosomal functions of RPs are numerous, and have been recently reviewed (Warner and McIntosh 2009; Zhou et al. 2015).

Given their role in regulating gene translation, cellular differentiation, and organismal development, it is perhaps unsurprising that altered RP expression has been implicated in human pathology. Indeed, an entire class of diseases has been shown to be associated with haploinsufficient expression or mutation in individual RPs. These so-called “ribosomopathies,” including Diamond-Blackfan Anemia (DBA) and Shwachman-Diamond Syndrome (SDS), are characterized by early onset bone marrow failure, variable developmental abnormalities and a life-long cancer predisposition that commonly involves non-hematopoietic tissues (Ruggero and Shimamura 2014; Yelick and Trainor 2015). The loss of proper RP stoichiometry and ensuing ribosomal stress result in increased ribosome-free RPs, which bind to MDM2 and impair its ubiquitin-mediated degradation of p53 (Boultwood et al. 2012; Shenoy et al. 2012; Ruggero and Shimamura 2014; Russo and Russo 2017). The resulting p53 stability is believed to underlie the bone marrow failure affecting erythroid or myeloid lineages in DBA and SDS, respectively. The developmental abnormalities of the ribosomopathies are variable and associate with specific RP loss or mutation. RPL5 loss in DBA, for example, is specifically associated with cleft palate and other craniofacial abnormalities whereas RPL11 loss is associated with isolated thumb malformations (Gazda et al. 2008).

Ribosomopathy-like properties have also been observed in various cancers. We have recently shown that RP transcripts (RPTs) were dysregulated in two murine models of hepatoblastoma and hepatocellular carcinoma in a tumor specific manner and in patterns unrelated to tumor growth rates (Kulkarni, under review). These murine tumors also displayed abnormal rRNA processing and increased binding of free RPs to MDM2, reminiscent of the aforementioned inherited ribosomopathies.

Perturbations of RP expression have been found in numerous human cancers, including those of the breast, pancreas, bladder, brain and many other tissues (Lai and Xu 2007; Artero-Castro et al. 2011; Jung et al. 2011; Hong et al. 2014; Kardos et al. 2014; Khan et al. 2015; Paquet et al. 2015; Yong et al. 2015; Russo et al. 2016; Sim et al. 2016; Fan et al. 2017; Russo et al. 2017; Shi et al. 2017). Mutations and deletions of RP-encoding genes have also been found in endometrial cancer, colorectal cancer, glioma, and various hematopoietic malignancies (Goudarzi and Lindstrom 2016; Ajore et al. 2017; Fancello et al. 2017). Indeed, the Chr. 5q- abnormality associated with myelodysplastic syndrome and the accompanying haploinsufficiency of RPS14 is considered one of the prototype “acquired” ribosomopathies that are often classified together with DBA, SDS and other inherited ribosomopathies (Ruggero and Shimamura 2014). Although many free RPs can induce cellular senescence during ribosomal stress via MDM2/p53, not all RPs possess such tumor suppressor functions; RPS3A, for example, transforms NIH3T3 mouse fibroblasts and induces tumor formation in nude mice (Naora et al. 1998).

A recent attempt to summarize the heterogeneity of RPT expression in human cancers was limited to describing expression differences of single RPTs among cancer cohorts, without accounting for larger patterns of variation that might better distinguish tumors from one another (Guimaraes and Zavolan 2016). RPT expression patterns were, however, examined in normal tissues using the dimensionality-reduction technique Principal Component Analysis (PCA) in the aforementioned study. These results showed hints of cell-specific patterning in the hematopoietic tissues examined, but not all cell types clustered into obviously distinct groups.

In the current work, we leverage a machine learning technique known as *t*-distributed stochastic neighbor embedding (*t*-SNE) to identify distinct patterns of RPT expression across both normal human tissues and cancers. Like PCA, *t*-SNE is a dimensionality reduction technique used to visualize patterns in a data set (van der Maaten 2008). With either technique, patterns shared between data points are represented with clustering. *t*-SNE differs from PCA in that it performs particularly well with highly dimensional data and is able to distinguish non-linear relationships and patterns. With this technique, we show that virtually all normal tissues and tumors can be reliably distinguished from one another based on their RPT expression profile. Tumors are readily distinguishable from normal tissues, but retain sufficient normal tissue patterning to allow for their origin to be easily discerned. Finally, a number of cancers possess subtypes of RPT expression patterns that correlate in readily understandable ways with molecular markers, various tumor phenotypes, and survival.

## Results

### t-SNE identifies tissue- and tumor-specific RPT expression

RNA-seq expression data for 9844 tumors (30 cancer types) and 716 matched normal tissues were obtained from The Cancer Genome Atlas (TCGA). Relative expression of RPTs was calculated for all samples and first analyzed using PCA. Normal tissue samples could, to a modest degree, be distinguished by their RPT expression patterns, though many tissue types demonstrated considerable overlap (**Figure 1A** and **Supplementary Fig. 1A**). Patterns of RPT expression in tumors were even more heterogeneous, and most cancer cohorts did not cluster discretely (**Figure 1B**).

**Figure 1.**
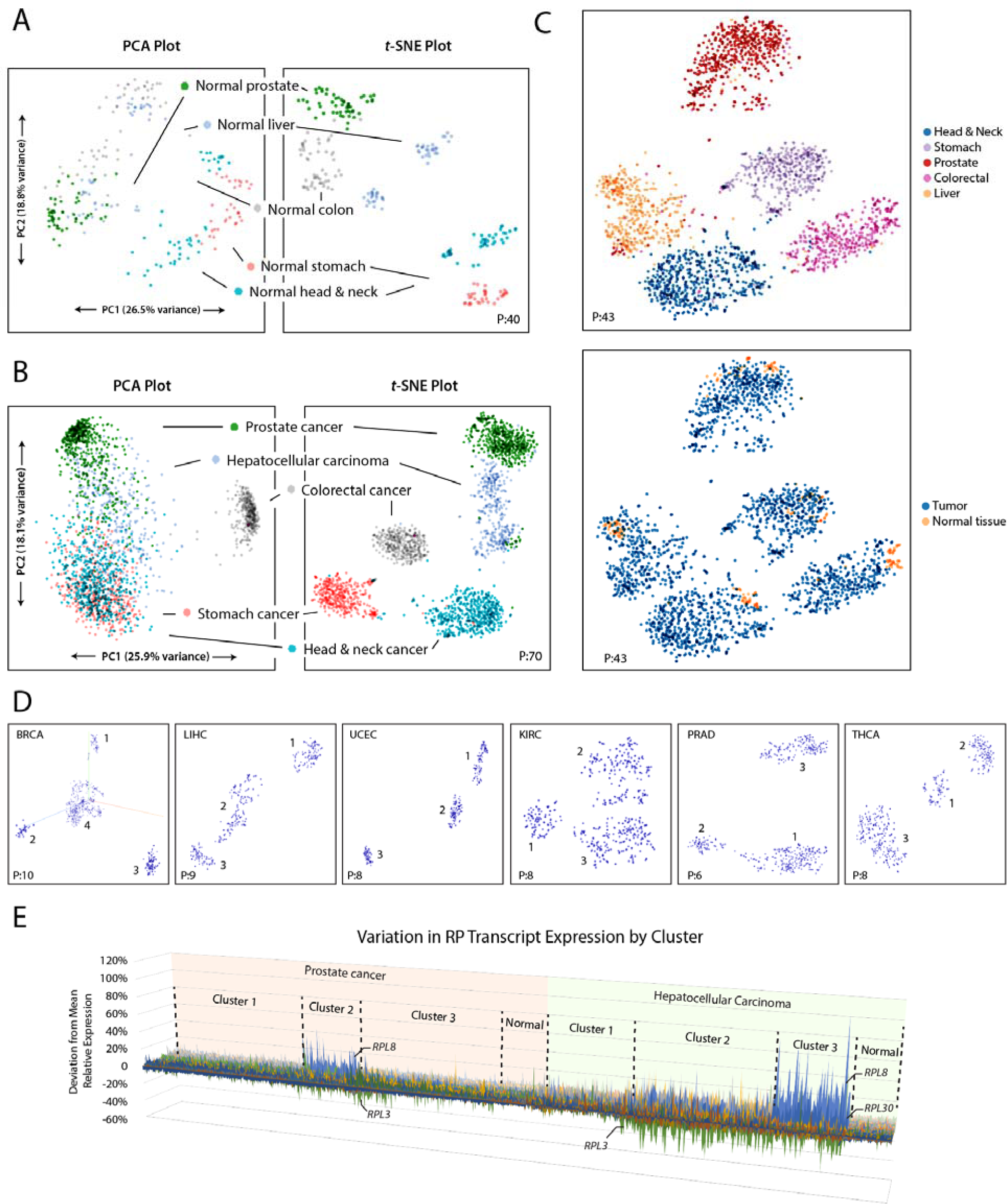
*t*-SNE better identifies clusters of RPT expression than PCA. A. Relative expression of RPTs in normal tissues from five cohorts was analyzed with PCA. In both methods, clustering occurs when samples possess similar underlying patterns of variation. *t*-SNE provides more distinct clusters that better associate with tissue of origin, indicating that normal tissues have distinct patterns of RPT expression. Axes are not labeled with *t*-SNE, as points are not mapped linearly and axes are not directly interpretable. **B**. Similar analyses in tumors. PCA clusters are poorly defined and do not correlate strongly with tumor type. *t*-SNE clusters are distinct and strongly associate with cancer type, indicating that tumors possess unique patterns of RPT expression based on their tissue of origin. **C**. Combined *t*-SNE analysis of RPT expression in normal tissue and tumor samples. Normal tissues and tumors cluster together but can be distinguished from one another, indicating that the latter retain a pattern of RPT expression resembling that of the normal tissue from which they originated. **D.** Many single cancer cohorts demonstrate sub-clustering by *t*-SNE. Clustering of six cohorts are provided as examples here. The number of clusters found in each cohort is listed in **Supplementary Table 1. E**. 3D area map of RPT relative expression in tumors from two cancer cohorts, sorted by cluster. The x-axis represents individual tumors, the z-axis represents individual RPTs, and the y-axis represents deviation from the mean relative expression. Cluster 2 of prostate cancer and Cluster 3 of HCC are both comprised of tumors with high relative expression of *RPL8* and low *RPL3*. See **Supplementary Figs. 1, 2**, and **5** for additional *t*-SNE plots of tumors and normal tissues. Perplexity settings for *t*-SNE analyses are designated in each plot by “P:”. For all analyses, learning rate (epsilon) = 10 and iterations = 5000.

Samples were then analyzed with *t*-SNE, which more clearly identified clusters of variation due to its ability to identify non-linear relationships between RPTs (**Figure 1A** and **B** and **Supplementary Fig. 1B**). Clustering of normal tissue samples correlated near perfectly with tissue type. Tumors also demonstrated clustering that strongly associated with tissue type, with 20 cohorts possessing largely distinct, non-overlapping clusters of tumors. When both normal tissues and tumors were analyzed together with *t*-SNE, samples also generally grouped into large clusters according to tissue type. Normal tissues, however, localized into smaller sub-clusters distinct from tumors (**Figure 1C** and **Supplementary Fig. 2**). Thus, while samples nearly always possessed RPT expression specific to their tissue type, normal tissues and tumors could be readily distinguished from one another.

Five cohorts – cholangiocarcinoma (CHOL), lung (LUNG), bladder (BLCA), cervical (CESC), and uterine carcinosarcoma (UCS) – were comprised of tumors that lacked tissue-specific RPT expression profiles and did not form distinct clusters. These tumors displayed significant overlap with each other as well as with tumors from the remaining five cohorts – liver (LIHC), colorectal (COADREAD), mesothelioma (MESO), pancreatic (PAAD), and skin cutaneous melanoma (SKCM) – which otherwise clustered distinctly from one another (**Supplementary Fig. 3**). Additionally, two clusters of tumors were found that did not associate with tissue of origin (**Supplementary Fig. 4**). The first contained 143 tumors from 15 cohorts, 98% of which had amplification and relative up-regulation of *RPL19, RPL23*, and *ERBB2* (Her2/Neu). The second contained 77 tumors from 12 cohorts with no discernable or unifying RPT expression pattern.

### t-SNE identifies sub-types of RPT expression within cancer types

Analyzed individually, 19 of 30 cancer types demonstrated sub-clustering of RPT expression with *t*-SNE (**Figure 1D, Supplementary Fig. 5**, and **Supplementary Table 1**). Graphing RPT relative expression by cluster using a 3D area map illustrated the different patterns of expression detected by *t*-SNE (**Figure 1E**). In some cases, these clusters differed from one another in the expression pattern of numerous RPTs, as with Clusters 1 and 3 of prostate cancer. In other cases, expression patterns appeared to be dominated by the differential relative expression of one or two RPTs, as with prostate cancer Cluster 2 and HCC Cluster 3, both of which possess tumors that overexpress *RPL8* and under-express *RPL3* (**Figure 1E**). While all clusters were distinct from normal tissues (**Figure 1C** and **Supplementary Fig. 2**), some clusters were more similar to normal tissues than others, such as prostate cancer Cluster 1 and HCC Cluster 1 (**Figure 1E**).

### Classification models

While *t*-SNE analyses are useful for visualization and pattern discovery, they do not alone provide a direct means for classification of future samples. Thus, with the knowledge that RPTs have both tissue- and tumor-specific expression patterns, we constructed various tumor classifier models based on these patterns. The constructed models consisted of both artificial neural network (ANN) and logistic regression (LR) classifiers, and are listed in **Supplementary Table 2**. An ANN model classified tumors by RPT content according to their tissue of origin on a separate test set with 93% accuracy. Similarly, a LR model distinguished tumors from normal tissues with >98% accuracy. Other LR models could distinguish glioblastoma multiforme tumors from other brain cancers with 100% accuracy and stratify both uterine and kidney clear cell tumors according to prognostic group with >95% accuracy.

### Characterizing tumor clusters identified by t-SNE

In order to quantify the differences in RPT expression that exist between clusters of tumors identified by *t*-SNE, RPT relative expression was compared between clusters of tumors with Analysis of Variance (ANOVA) and graphed with volcano plots (Figures 2 and 3A). Small but highly significant differences in the expression of dozens of RPTs occurred in nearly every tumor cluster (P as low as 10^-220^). As was the case with prostate cancer and HCC, expression patterns in clusters were often dominated by particularly significant differences in expression of one or two RPTs, most commonly *RPL3, RPS4X, RPL8, RPL30*, and *RPL13*. Other tumor clusters, notably those involving the uterus, brain, and lung, possessed more complex differences involving many RPTs (Figures 2 and 3A).

**Figure 2.**
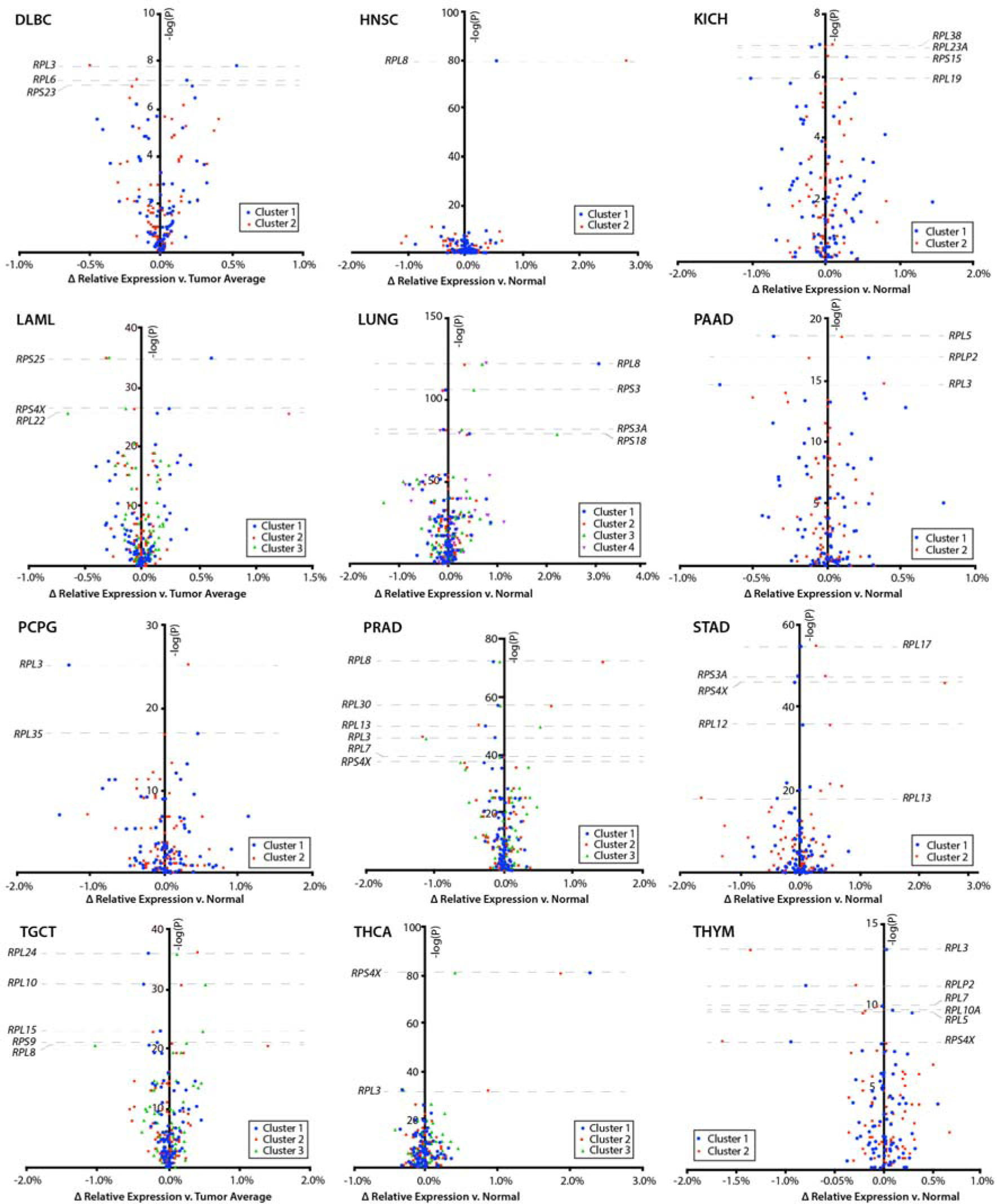
Volcano plots of relative RPT expression in tumor clusters in twelve cancer cohorts. Relative expression of RPTs was compared between tumor clusters in each included cancer cohort with ANOVA tests. The negative log of the ANOVA P-value for each RPT is displayed on the y-axis and the difference in relative expression across tumor clusters is displayed on the x-axis. RPTs near the top of the graphs are most significantly differentially expressed between tumor clusters. Note that nearly every RPT in virtually all cancer cohorts falls above –log(P) of 2, corresponding to P < 0.01 and indicating that tumor clusters have significantly distinct expression of virtually all RPTs. For each cohort, the number of samples in each cluster are shown under the label “*n*”. Additional volcano plots of seven other cancer cohorts are continued in **Figure 3**.

**Figure 3.**
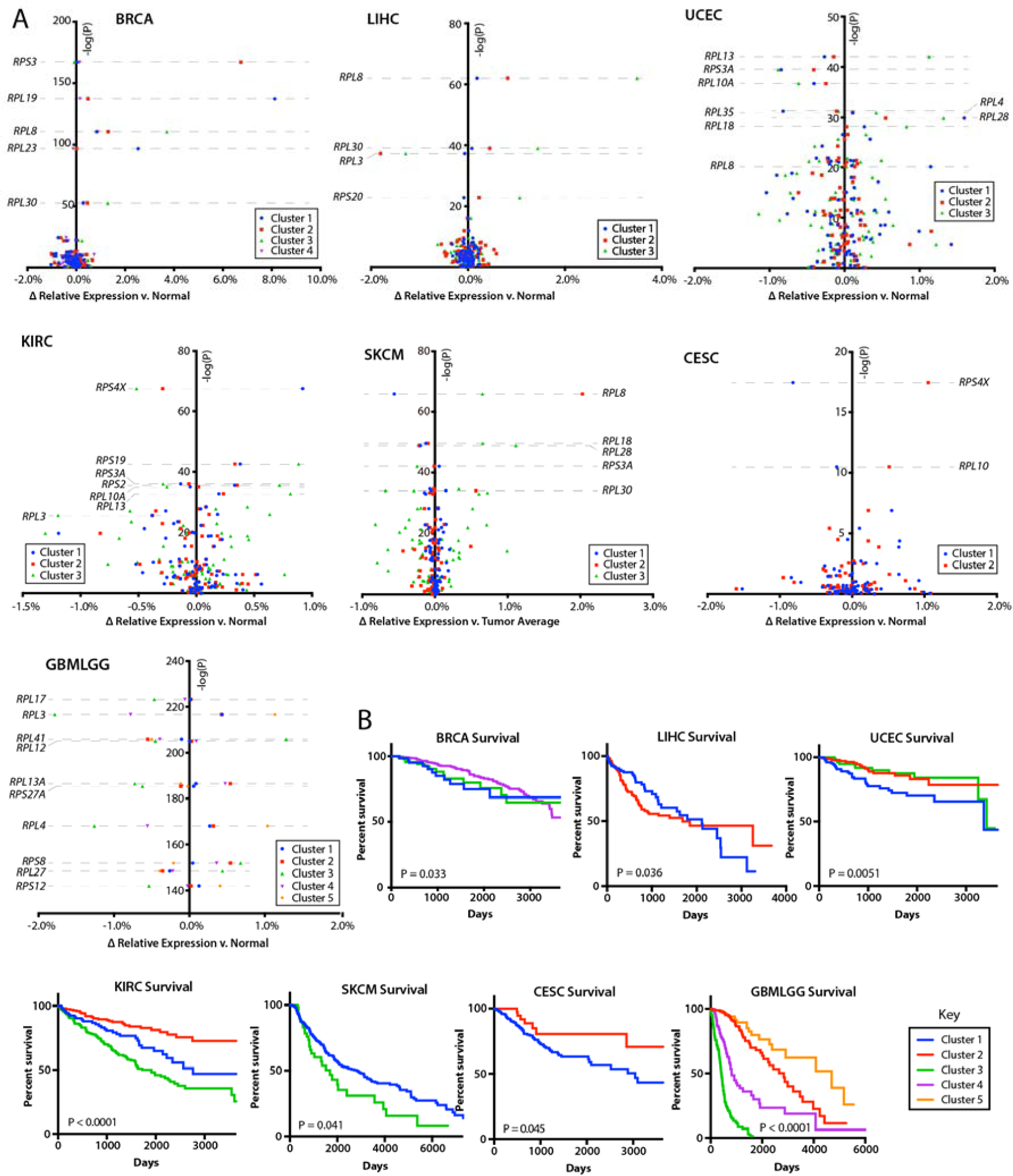
Volcano plots of relative RPT expression in tumor clusters associated with survival. A. Volcano plots comparing RPT relative expression between tumor clusters were generated, as in Figure 2, for the remaining seven cancer cohorts which possessed tumor sub-clustering by *t*-SNE. Note that for the sake of clarity, clusters 5 and 6 are excluded from the LUNG cohort plot. These clusters correlated near perfectly with amplification and highly significant up-regulation of *RPS3* and *RPS16*, respectively (**Table 2**). **B.** Patient survival by *t*-SNE cluster. Of the 19 cancer cohorts with sub-clustering of RPT expression patterns by *t*-SNE, seven possessed clusters that correlated with survival. Significance was determined with log-rank and Wilcoxon rank sum tests where appropriate, using all survival data available, including any data points beyond what are displayed in the survival curves.

Several recurrent alterations in RPT expression were found among the 19 cancer cohorts with sub-clustering (Table 1). Nine of these clusters, arising from thyroid, brain, liver, kidney clear cell, thymoma, prostate, pancreatic, pheochromocytoma and paraganglioma, and B-cell lymphoma, contained tumors with low relative expression of *RPL3*. These clusters also shared expression patterns with other RPTs, including the relative down-regulation of *RPL5* and up-regulation of *RPL36* and *RPL38*. Excluding thymomas, all other tumor clusters with low *RPL3* also shared 11 other similarly co-regulated RPTs. Additionally, six cancer cohorts – prostate, breast, liver, lung, melanoma, and head and neck – contained tumor clusters distinguished by overexpression of *RPL8, RPL30* and *RPS20*, with shared expression patterns of 19 other RPTs. Relative up-regulation of *RPS4X* occurred in tumors from six cohorts, all of which showed similar co-expression patterns of nine other RPTs. Finally, tumor clusters overexpressing *RPL13* were found in prostate, uterine and kidney clear cell carcinoma and shared similar patterns of expression of 42 other RPTs (Figures 2 and 3A and Table 1).

**Table 1.** Recurring patterns of RPT relative expression across cancer cohorts. Certain patterns of expression distinguishing tumor clusters from one another were observed in multiple clusters across cancer cohorts, as shown in Figure 2 and 3A. In this table, “low” refers to tumor clusters expressing lower relative expression of a given RPT relative to other tumors in the given cancer cohort, and “high” refers to clusters with greater relative expression compared to other tumors.

| Observed pattern | Cohort | Cluster | Co-regulated RPTs |  |
| --- | --- | --- | --- | --- |
|  |  |  | Lower expression | Higher expression |
| Low *RPL3* | THCA | 1 |  |  |
|  | GBMLGG | 3, 4 | *RPL5, RPL10, RPL17,* | *RPL28, RPL35, RPL36, RPL38,* |
|  | LIHC | 2, 3 | *RPL22, RPL26, RPS23* | *RPLP2* |
|  | KIRC | 3 |  |  |
|  | THYM* | 2 |  |  |
|  | PRAD | 2, 3 |  |  |
|  | PAAD | 1 |  |  |
|  | PCPG | 1 |  |  |
|  | DLBC | 2 |  |  |
| High *RPL8* and *RPL30* | BRCA | 3 | *RPL10, RPL13A, RPL19,* | *RPL36A, RPL7, RPS20* |
|  | LIHC | 3 | *RPL21, RPL34, RPL37A,* |  |
|  | PRAD | 2 | *RPL4, UBA52, RPL5, RPS11,* |  |
|  | LUNG | 1 | *RPS12, RPS13, RPS17,* |  |
|  | SKCM | 2 | *RPS27A, RPS8, RPS9* |  |
|  | HNSC | 2 |  |  |
| High *RPS4X* | PRAD | 1 | *RPL13, RPL23, RPL37,* | *RPS3A, RPL17* |
|  | KIRC | 1 | *RPL8, RPLP1, RPS16, RPS2* |  |
|  | THCA | 1, 2 |  |  |
|  | STAD | 2 |  |  |
|  | LAML | 1 |  |  |
|  | CESC | 2 |  |  |
| High *RPL13* | PRAD | 3 | *RPL10, RPL10A, RPL26,* | *RPL12, RPL13, RPL18, RPL24,* |
|  | UCEC | 3 | *RPL3, RPL35A, RPL41,* | *RPL27A, RPL28, RPL32,* |
|  | KIRC | 3 | *RPL5, RPL7, RPL7A, RPS12,* | *RPL35, RPL36, RPL37, RPL38,* |
|  |  |  | *RPS13, RPS14, RPS15A,* | *RPL39, RPLP1, RPLP2, RPS15,* |
|  |  |  | *RPS18, RPS23, RPS27, RPS3,* | *RPS17, RPS19, RPS20, RPS24,* |
|  |  |  | *RPS3A, RPS4X, RPS5* | *FAU, RPS4Y1, RPS9* |

\* Although the main observed pattern was found in this tumor cluster, other RPTs were not co-regulated in the same manner

In some cases, RP gene copy number variations (CNVs) were associated with clustering (**Table 2**). Notably, the aforementioned *RPL8/RPL30* overexpression pattern strongly correlated with co-amplification of a region on 8q22-24 containing *RPL8, RPL30*, and *MYC*. Similarly, an amplicon containing *RPL19, RPL23*, and *ERBB2* (Her2/Neu) was amplified in 99% of the breast cancers in Cluster 1. Some tumor clusters associated with specific CNVs to a lesser degree. For example, 48% of tumors in kidney clear cell carcinoma Cluster 3 possessed deletions of *RPL12, RPL35*, and *RPL7A* on 9q33-34. Similarly, half of brain cancers in Cluster 1 possessed a 1p/19q13 co-deletion, compared to nearly 100% of tumors in Cluster 5 with this deletion (**Table 2**). Other tumor clusters in various cancer cohorts had differences in overall CNV frequencies. In testicular cancer, 39 RP genes were amplified at different frequencies among the three clusters. Endometrial cancer Cluster 1 and HCC Cluster 2 had more CNVs overall, but no RP gene was amplified or deleted with a frequency of greater than 65% in any given tumor cluster.

**Table 2.**
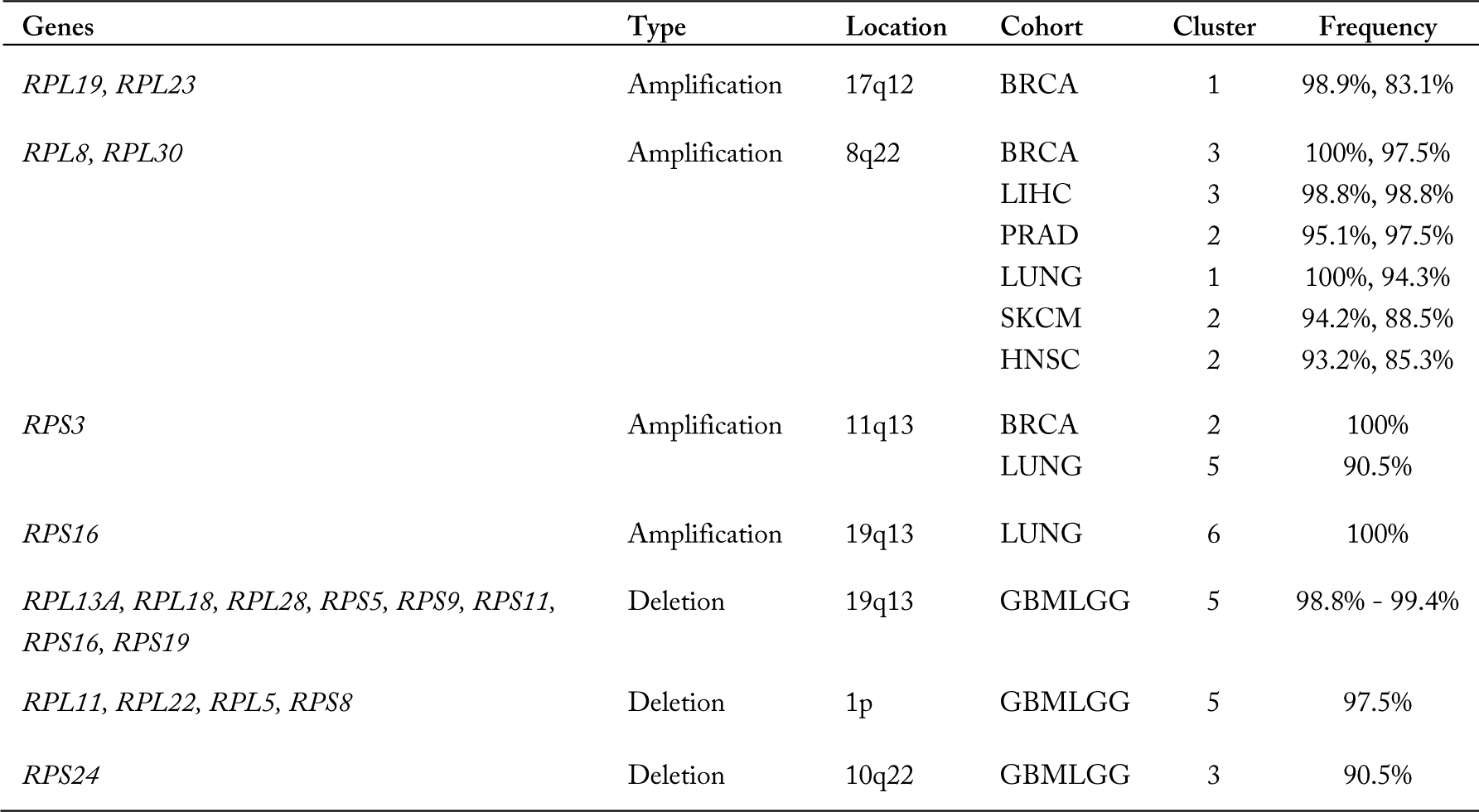
RP gene copy number alterations associated with *t*-SNE clusters. Some tumor clusters were significantly associated with greater incidence of copy number alterations than other tumors from the same cancer cohorts (α < 0.01); clusters with >90% of tumors possessing a given copy number alteration are included in this table.

Many tumor clusters – each representing a distinct RPT expression pattern - significantly associated with various clinical parameters, molecular markers, and tumor phenotypes (**Table 3**). This was particularly true for brain cancer, testicular cancer, thyroid cancer, lung cancer, and endometrial cancer. Tumor clusters in HCC and head and neck cancers strongly correlated with etiologically-linked infections. For example, chronic hepatitis B infection was 2-fold more common in HCC patients with Cluster 2 tumors compared to other HCC patients. Similarly, chronic HPV infection was 4.7-fold more frequent in head and neck cancer patients with Cluster 1 tumors compared to other patients in this cohort. Patient gender also associated with tumor clustering to varying but significant degrees in kidney clear cell carcinoma and AML. Notably, these clusters also associated with differential relative expression of the X-chromosome encoded *RPS4X*. Other clinical markers and tumor phenotypes significantly associated with tumor clustering can be found in **Table 3**.

**Table 3.** Tumor phenotypes and clinical parameters associated with *t*-SNE clustering. Tumor phenotypes and clinical markers were compared between tumor clusters using Chi-squared tests, with significance defined as α < 0.01. “Other tumors” are comprised of all tumors from the same cancer cohort not falling into the given cluster. Data were obtained using the UC Santa Cruz Xenabrowser, under the data heading “Phenotypes.”

| Feature | Cluster | Comparison | P-value |
| --- | --- | --- | --- |
| *Breast cancer* |  |  |  |
| Her2/Neu status | Cluster 1 – 88.2% | Other tumors – 7.7% | < 0.0001 |
| *Hepatocellular carcinoma* |  |  |  |
| Hepatitis B | Cluster 2 – 38.4% | Other tumors – 18.6% | < 0.0001 |
| *Endometrial carcinoma* |  |  |  |
| Type II (serous) type | Cluster 1 – 32.4% | Other tumors – 5.2% | < 0.0001 |
| Locoregional disease or recurrence | Cluster 1 – 8.6% | Other tumors – 2.2% | 0.004 |
| *Lung cancer* |  |  |  |
| Adenocarcinoma | Cluster 2 – 80.0% | Other tumors – 37.8% | < 0.0001 |
| Squamous cell carcinoma | Cluster 4 – 73.6% | Other tumors – 34.2% | < 0.0001 |
| *Kidney clear cell carcinoma* |  |  |  |
| Gender | Cluster 1 – 92.9% female | Other tumors – 29.0% female |  |
| Longest dimension | Cluster 2 – 1.5 cm | Other tumors – 1.8 cm | < 0.0001 |
| Histologic grade 3 or 4 | Cluster 3 – 69.0% | Other tumors – 41.5% | < 0.0001 |
| Pathologic stage III or IV | Cluster 3 – 58.5% | Other tumors – 25.5% | < 0.0001 |
| *Thyroid carcinoma* |  |  |  |
| Gender | Cluster 3 – 35.4% male | Other tumors – 11.3% male | < 0.0001 |
| Tall cell subtype | Cluster 1 – 17.2% | Other tumors – 5.03% | 0.0001 |
| Follicular subtype | Cluster 3 – 27.7% | Other tumors – 10.5% | < 0.0001 |
| *Acute myeloid leukemia* |  |  |  |
| Gender | Cluster 1 – 100% female<br>Cluster 3 – 81.5% male | Cluster 2 – 53.6% female | < 0.0001 |
| Complex (>3 distinct) cytogenetic abnormalities | Cluster 3 – 21.0% | Other tumors – 5.5% | 0.0025 |
| *Testicular cancer* |  |  |  |
| Seminoma subtype | Cluster 1 – 100% | Other tumors – 14.5% | < 0.0001 |
| Embryonal carcinoma subtype | Cluster 2 – 94.6% | Other tumors – 20.6% | < 0.0001 |
| Teratoma subtype | Cluster 3 – 84.0% | Other tumors – 14.7% | < 0.0001 |
| *Head and neck cancer* |  |  |  |
| HPV infection | Cluster 1 – 46.1% | Other tumors – 9.8% | 0.0001 |
| *Skin cutaneous melanoma* |  |  |  |
| Breslow depth | Cluster 1 – 4.8 cm | Other tumors – 7.6 cm | 0.0083 |
| *Brain cancer* |  |  |  |
| Glioblastoma | Cluster 3 – 100% | Other tumors – 0% | < 0.0001 |
| Astrocytoma low-grade glioma | Cluster 2 – 53.0%<br>Cluster 4 – 62.7% | Other tumors – 2.5% | < 0.0001 |
| Non-astrocytoma low-grade glioma | Cluster 1 – 86.7%<br>Cluster 5 – 97.0% | Other tumors – 29.1% | < 0.0001 |

Tumor clusters were often predictive of survival, including some clusters that did not significantly associate with any other known tumor subtype (**Figure 3B**). For example, Clusters 2 and 4 of the brain cancer cohort, which could not otherwise be distinguished by any known clinical parameter or tumor subtype, possessed vastly different survival patterns. Other cancer cohorts with significant survival differences among clusters included breast, liver, endometrial, kidney clear cell, melanoma, and cervical cancers.

## Discussion

By investigating expression patterns of individual RPTs and utilizing more traditional and less powerful linear forms of dimensionality reduction such as PCA, previous studies have found modest evidence of tissue-specific patterning of RPT expression in some normal tissues (Guimaraes and Zavolan 2016). Extending these types of analyses to tumors has been largely unfruitful, presumably due to the complex regulation of RPT expression and because many of the RPT relationships are non-linear. As shown here, however, the machine learning algorithm *t*-SNE provides a more elegant and robust dimensionality reduction that better highlights distinct patterns of RPT expression in both tumors and the normal tissues from which they arise.

Consistent with more restricted and tentative conclusions of previous findings, our results using *t*-SNE clearly demonstrate that RPT expression patterns are not only tissue-specific but provide the ability to define tissue and tumor differences with a heretofore unachievable degree of resolution. The small cluster of 77 neoplasms that did not associate with their respective tissue clusters (**Supplementary Fig. 4**) may represent either a subset of tumors that have lost control of their underlying tissue-specific expression patterns or that originated from a minority subpopulation of normal cells whose RPT expression is not representative of the remainder of the tissue.

In addition to their tissue-specific patterning, virtually all tumors showed perturbations of RPT expression that readily allowed them to be distinguished from normal tissues. In some cancers, the tumor-specific patterning of RPT expression was relatively homogeneous and could not otherwise be subcategorized. Most cohorts, however, were comprised of subgroups of tumors with distinct RPT expression patterns, all of which remained distinguishable from normal tissue. The fact that many of these patterns correlated with molecular and clinical features implicates RPT expression patterns in tumor biology.

Aside from potentially altering translation, the notion that altered RP expression might influence the behaviors of both normal tissues and tumors is not new. In the ribosomopathies, the binding of any one of about a dozen RPs to MDM2 with subsequent stabilization of p53 is thought to underlie bone marrow failure (Boultwood et al. 2012; Shenoy et al. 2012; Ruggero and Shimamura 2014). It has been proposed that subsequent circumvention of this p53-mediated senescence by mutation and/or dysregulation of the p19^*ARF*^/MDM2/p53 pathway is responsible for the propensity for eventual neoplastic progression (De Keersmaecker 2015). In cancers, the binding of free RPs to MDM2 has been shown to mediate the response to ribosomal-stress-inducing chemotherapeutics such as actinomycin D and 5-fluorouracil (Sun et al. 2007; Esposito et al. 2014; Russo et al. 2016).

Individual RPs have also been associated with specific tumor phenotypes. For example, RPL3 regulates chemotherapy response in certain lung and colon cancers, associates with the high-risk neuroblastoma subtype, and may have a role in the acquisition of lung cancer multidrug resistance (Khan et al. 2015; Russo et al. 2016; Russo et al. 2017). Breast cancers with elevated expression of *RPL19* are more sensitive to apoptosis mediated drugs that induce endoplasmic reticulum stress (Hong et al. 2014). *RPS11* and *RPS20* have been proposed as prognostic markers in glioblastoma (Yong et al. 2015) and the down-regulation of *RPL10* correlates with altered treatment response to dimethylaminoparthenolide (DMAPT) in pancreatic cancer (Shi et al. 2017).

Our results also extend the findings of previous studies by demonstrating that in the vast majority of cancers, subsets of RPTs are expressed coordinately and have additional interpretive power when examined in the context of global RPT expression patterning. This suggests that further insights into the roles RPTs have in tumor development may be revealed by evaluating RPT relative expression. For example, the regulation of chemotherapy response by RPL3 may be found to occur in other cancer types once the expression of *RPL3* relative to other RPTs has been taken into account. The apparent crucial role of RPT patterning in tumors may explain why a previous study found conflicting results when examining the expression of individual RPs in tumors (Lai and Xu 2007).

Our results suggest a more ubiquitous role for RPL3 in regulating tumor phenotypes, beyond that already described in colorectal carcinoma, lung cancers, and neuroblastoma (Khan et al. 2015; Russo et al. 2016; Russo et al. 2017). Of the recurring RPT expression patterns discovered by *t*-SNE, the pattern associated with *RPL3* down-regulation occurred most frequently, involving tumors from nine cancer cohorts. Many clusters of tumors with down-regulated *RPL3* possessed inferior survival, including those from liver, kidney clear cell, and brain cancers. The fact that relative down-regulation of *RPL3* occurred in these tumor clusters with predictable expression of 11 other RPTs suggests that RPL3 may be acting in concert with these other identified RPs to exert its effects.

Other recurring patterns of RPT expression across cancer cohorts involved *RPS4X, RPL13, RPL8* and *RPL30* (**Table 1**). Altered *RPS4X* expression, found in six cancer cohorts, associated with unique expression of nine other RPTs, strongly suggesting an underlying coordinated expression. As with *RPL3*, deregulated *RPS4X* has been previously associated with various tumors and tumor phenotypes, including subgroups of colorectal carcinoma, a myelodysplasia risk signature and poor prognosis in bladder cancer (Sridhar et al. 2009; Jung et al. 2011; Paquet et al. 2015). Interestingly, some of our tumor clusters with altered *RPS4X* expression were comprised of a greater proportion of females than males (**Table 1** and **Table 3**), perhaps reflecting the fact that the *RPS4X* gene resides on chromosome X. Although the cause of perturbed *RPS4X* expression in these tumor clusters is unknown, altered methylation patterns on chromosome X have been described in different subsets of cancers (Spatz et al. 2004; Chaligné et al. 2015) and could be responsible for the *RPS4X* expression patterns detected by *t*-SNE.

Unlike *RPL3* and *RPS4X*, the role of *RPL13* in tumor development is less clear. Activation of RPL13 has been described in a subset of gastrointestinal malignancies and correlated with greater proliferative capacity and attenuated chemoresistance (Kobayashi et al. 2006), but further evidence for a role of RPL13 in tumor development is lacking. Furthermore, clinical correlations of the prostate, uterine and kidney cancer *t*-SNE clusters described here with relative overexpression of *RPL13* were inconsistent. Uterine cancers with high relative *RPL13* tended to correlate with favorable survival, whereas prostate cancers with high *RPL13* showed no differences in prognosis or clinical features, and kidney clear cell carcinomas with high *RPL13* tended to be of higher pathologic grade and conferred significantly poorer survival (**Table 1, Table 3**, and **Figure 3B**). The fact that these clusters shared similar patterning of 42 other RPTs, however, suggests that the inciting factors responsible for higher *RPL13* expression are not only shared by these tumors but coordinately regulate a common subset of RPTs.

In some cases, RPT expression patterns could be accounted for in part by CNVs, as exemplified by the recurrent *RPL8* and *RPL30* overexpression pattern (**Tables 1** and **2**). Virtually all tumors with this expression pattern possessed co-amplification of a region on 8q22-24 that includes *RPL8, RPL30* and the oncogene *MYC*. Amplification of this region has been previously described in breast cancers and correlates with chemoresistance and metastasis (Hu et al. 2009; Parris et al. 2014; Taghavi et al. 2016). Our results indicate that this amplification and the ensuing overexpression of *RPL8* and *RPL30* also occurs in subsets of melanoma, liver, prostate, lung, and head and neck cancers. CNVs in *RPL19* and *RPL23* in breast cancer (**Table 2**) likely occur due to their co-amplification with *ERBB2* on 17q12. Overexpression of *RPL19* has previously been described in a subset of breast cancers (Hong et al. 2014). The small cluster of 144 tumors that did not group according to tissue of origin (**Supplementary Fig. 4**), comprised of tumors from 15 cohorts, also shared amplification of this region on 17q12, indicating that this CNV is not restricted to breast cancers and ultimately affects global RPT expression patterning. Amplification of a region on 11q13 that contains *RPS3*, occurring in a cluster of breast cancers and HCCs, has been previously described in both cancers and is thought to confer unfavorable prognosis due to amplification of the oncogene *EMS1* in this region (Ormandy et al. 2003; Yuan et al. 2003). The co-deletion of 19q13 with regions of 1p, which include numerous RP genes, has been described in low-grade gliomas and correlates with a favorable prognosis (Barbashina et al. 2005; Vogazianou et al. 2010).

The co-overexpression *RPS25* and *RPS4X* detected in one cluster of AML (**Figure 2**) has been previously identified as contributing to the poor risk signature in myelodysplastic syndrome (Sridhar et al. 2009). This also associated with significant differential expression of 37 RPTs, which is consistent with our finding that *RPS25* and *RPS4X* overexpression occur within the context of a larger and coordinated pattern of RPT expression. The *RPS25* and *RPS4X* overexpressing AML cases likely possess a similar molecular alteration to those with the poor risk signature in MDS.

Collectively, our findings provide strong evidence to support the notion that RPT regulation by both tumors and normal tissues is complex, ordered, and highly coordinated. Although the means by which altered RPT patterns influence the pathogenesis and/or behavior of tumors remain incompletely understood, several non-mutually exclusive mechanisms can be envisioned. First, changes in RP levels may influence overall ribosome composition, affecting the affinity for certain classes of transcripts and/or the efficiency with which they are translated. One such class of transcripts may be those with IRES elements, *cis*-regulatory sequences found in the 5’-untranslated regions of more than 10% of cellular mRNAs. IRES elements are found with particularly high frequency on transcripts encoding proteins involved in cell cycle control and various stress responses. Efficient translation of these IRES-containing transcripts has been shown to depend on the presence of specific RPs, notably *RPS25, RPS19* and *RPL11* (Landry et al. 2009; Muhs et al. 2011; Horos et al. 2012). Changes in ribosome affinity for IRES elements have been shown to reduce translation of tumor suppressors such as p27 and p53 and to promote cancer development (Bellodi et al. 2010).

RPs may also influence cancer development via extra-ribosomal pathways. In addition to their promotion of p53 stability mediated by binding to and inactivating MDM2, specific RPs have been shown to inactivate Myc; to inhibit the Myc target Lin28B; to activate NF-κB, cyclins, and cyclin-dependent kinases and to regulate a variety of other tumorigenic functions and immunogenic pathways (Warner and McIntosh 2009; Zhou et al. 2015).

In addition to providing evidence that tumors may use RPs to direct tumor phenotypes, our findings have allowed us to leverage the tissue- and tumor-specificity of RPT expression to generate highly sensitive and specific models that allow for precise tumor identification and sub-classification (**Supplementary Table 2**). Clinically, these might be useful for determining the tissue of origin of undifferentiated tumors and for predicting long-term behaviors in otherwise homogeneous cancers such as in kidney clear cell carcinoma and those of the central nervous system (**Figure 3B**). With more samples and further refinement to ANN structures, future iterations of these models will likely have even greater discriminatory power.

A limitation of using data from TCGA is the fact that transcript expression does not always correlate with protein expression, particularly in cancers (Chen et al. 2002; Tian et al. 2004; Koussounadis et al. 2015). Thus, it is difficult to predict how the different tissue-specific RPT expression patterns we identified correlate with actual protein expression in these cancers and/or with the numerous post-translational modifications that can alter RP behaviors. As this is a cross-sectional study, we also recognize that causality cannot be inferred and it remains unknown whether altered RPT expression is an early or late event in tumorigenesis despite its predictive value. Further molecular analyses of the identified *t*-SNE clusters with whole-transcriptome sequencing data, pathway analysis, whole-genome DNA mutation data, and DNA methylation patterning may offer additional insights into the biological mechanisms that link altered RPT expression with tumor phenotypes.

In summary, machine learning-based approaches have allowed us to determine that RPTs are expressed with distinct patterning across tissue types. This tissue-specificity persists in tumors, yet normal tissues and tumors can be readily distinguished from one another with high accuracy and confidence. Many cancers can be further sub-categorized into heretofore unrecognized, yet clinically important, subtypes based only upon RPT expression patterns. Several patterns of RPT expression recur across cancer types, suggesting common underlying and regulated modes of transcriptional regulation. Our results indicate that the expression of RPTs in tumors is biologically coordinated, clinically meaningful, and can be leveraged to create potential clinical tools for tumor classification and therapeutic stratification.

## Methods

### Accessing Ribosomal Protein Transcript Expression Data

RNA-seq whole-transcriptome expression data for 9844 tumors and 716 normal tissues from The Cancer Genome Atlas (TCGA) was accessed using the University of California Santa Cruz (UCSC) Xenabrowser (https://xenabrowser.net). Only primary tumors were included for analysis, apart from the melanoma (SKCM) cohort, as the vast majority of tumors with sequencing data in this cohort were metastatic (78%). For each of the 30 cancer cohorts, RNA-seq data was selected according to the label “gene expression RNAseq (polyA+ IlluminaHiSeq).” “IlluminaGA” RNA-seq expression data was used for the cohort Uterine Corpus Endometrial Carcinoma (UCEC), as this group of data had more samples than the “IlluminaHiSeq” group. For all cancer cohorts, expression data for 80 cytoplasmic RP genes were extracted and base-two exponentiated, as the raw RPKM (Reads Per Kilobase per Million mapped reads) expression data was stored log-transformed. The sum of total RPKM counts for all ribosomal protein genes were calculated for each sample, and relative expression of each RP gene in a sample was calculated by dividing the RPKM gene expression by this summed expression.

### Visualizing Ribosomal Protein Transcript Expression

Principal component analyses and *t*-SNE analyses of RPT relative expression in normal tissues and tumor samples were performed using TensorFlow r1.0 and Tensorboard (https://tensorflow.org). *t*-SNE analyses were performed at a learning rate (epsilon) of 10 with 5000 iterations or until the visualization stabilized. *t*-SNE was initially performed in two dimensions for all analyses; data sets that could not be cleanly visualized with two dimensions, particularly those with a large number of samples, were visualized with three-dimensional *t*-SNE. Multiple analyses were performed with perplexity settings varying between 6-15 for all individual cohort analyses and 10-30 for all grouped cohort analyses, with final perplexity settings for each analysis chosen to maximize cluster distinctions. Clusters of at least 10 samples which distinctly separated visually from other samples were named and samples from these clusters were identified. 3D area maps of RPT relative expression were generated using Microsoft Excel, with each sample listed across the x-axis, RPTs listed across the z-axis, and relative expression of each RPT across the y-axis.

### *Comparing* t*-SNE Clusters*

Relative expression of RPTs were compared between *t*-SNE clusters with Analyses of Variance (ANOVA) using R version 3.3.2 (http://www.R-project.org/). ANOVA p-values were log10-transformed and used to generate Volcano plots comparing expression patterns between clusters. Volcano plots were graphed with Graphpad Prism 7 (GraphPad Software, Inc., La Jolla, CA).

Clinical and survival data for each TCGA cancer cohort were accessed again using the UCSC Xenabrowser under the data heading “Phenotypes.” For each cohort, survival curves of tumors in each *t*-SNE cluster were compared with Mantel-Haenszel (log-rank) and Gehan-Breslow-Wilcoxon methods using Graphpad Prism 7. Categorical clinical variables were compared between clusters of tumors with Chi-squared tests. Continuous variables which were normally distributed were compared with *t*-tests assuming heteroskedasticity, and non-normally-distributed variables were compared with Wilcoxon sign-rank tests. All statistical tests were two-tailed.

### Co-regulated RPTs

Certain groups of RPTs possessed recurring, highly-significant differences between multiple *t*-SNE clusters, including *RPL3, RPL8, RPS4X*, and *RPL13*. For each TCGA cohort with a cluster that possessed significantly different relative expression of one of these transcripts, relative expression of all other RPTs was compared between the identified cluster and other tumors in the same cohort. Co-regulated transcripts were defined as those with consistent differences in relative expression when comparing clusters of interest to other tumors from the same cohort (**Table 1**). For example, five TCGA cohorts had a *t*-SNE cluster with significant relative overexpression of *RPL8* and *RPL30*. When comparing relative expression of other RPTs between these clusters and other tumors from the same cohorts, all five clusters with high *RPL8* and *RPL30* also displayed, on average, lower relative expression of *RPL10* and higher relative expression of *RPL7*.

### Ribosomal Protein Gene Copy Number Variations (CNVs)

CNV data for TCGA tumors was accessed using the UCSC Xenabrowser under the data heading “copy number (gistic2_thresholded).” Positive values were classified as amplifications, and negative values were classified as deletions. The frequency of amplifications and deletions in RP genes were compared between clusters of tumors in each TCGA cohort using Chi-squared tests and adjusted for 5% false discovery rate. Within each cancer cohort, clusters of tumors with significantly greater incidence of a CNV compared to other tumor clusters, and which possessed >90% incidence of this copy number variation, were included in **Table 2**.

### Classification Models

Using RPT relative expression in tumors and normal tissues, classification models were created using both logistic regression (LR) and feed-forward, fully-connected artificial neural networks (ANNs) (Dreiseitl and Ohno-Machado 2002). LR models were used for binary classifiers and developed with Stata SE 14 (StataCorp LP, College Station, TX) with *c*-statistics, sensitivity, and specificity reported in **Supplementary Table 2**. ANN models were generated for classifiers with multiple outcomes (*e.g*. tissue of origin models) and binary classifiers with a LR model that failed to converge.

ANN models were created and tested using TensorFlow with graphics processing unit (GPU) acceleration on a Titan X Pascal. To reduce bias, samples were balanced for both training and testing by cancer cohort such that each training and test set had the same number of samples from each cohort. 60% of data sets were used for training and 10% for validation and hyper-parameter tuning. Hyper-parameter sweeps were used to test all possible combinations of the following: learning rate (0.001, 0.002, 0.005, 0.01), batch size (100, 500, none), dropout rate (0.9, 0.95, 1), hidden layer structure (both one and two layers with sizes varying between 0 – 200 in increments of 25), and L2 regularization rate (0.00001, 0.0001, 0.001). All ANNs utilized ReLU activation functions. Neural network training performance was monitored with Tensorboard and stopped once validation accuracy had plateaued. The remaining 30% of data comprised a separate test set, which was used to test the final model’s classification accuracy once the hyper-parameters were chosen and the model trained. Performance of ANN models on the separate test sets were reported as classification accuracies in **Supplementary Table 2**.

## Data access

RNA-based sequencing data from this study was obtained from The Cancer Genome Atlas (TCGA; https://cancergenome.nih.gov/).

## Acknowledgements

This work was supported by NIH grant RO1 CA174713 and by The Hyundai Hope on Wheels Foundation to EVP. JMD was supported by the Clinical Scientist Training Program at the University of Pittsburgh School of Medicine. We also gratefully acknowledge the support of NVIDIA Corporation with the donation of the Titan X Pascal GPU used for this research.

## References

Ajore R, Raiser D, McConkey M, Joud M, Boidol B, Mar B, Saksena G, Weinstock DM, Armstrong S, Ellis SR et al. 2017. Deletion of ribosomal protein genes is a common vulnerability in human cancer, especially in concert with TP53 mutations. EMBO molecular medicine 9: 498–507.

Artero-Castro A, Castellvi J, Garcia A, Hernandez J, Ramon y Cajal S, Lleonart ME. 2011. Expression of the ribosomal proteins Rplp0, Rplp1, and Rplp2 in gynecologic tumors. Human pathology 42: 194–203.

Barbashina V, Salazar P, Holland EC, Rosenblum MK, Ladanyi M. 2005. Allelic losses at 1p36 and 19q13 in gliomas: correlation with histologic classification, definition of a 150-kb minimal deleted region on 1p36, and evaluation of CAMTA1 as a candidate tumor suppressor gene. Clinical cancer research: an official journal of the American Association for Cancer Research 11: 1119–1128.

Bellodi C, Krasnykh O, Haynes N, Theodoropoulou M, Peng G, Montanaro L, Ruggero D. 2010. Loss of function of the tumor suppressor DKC1 perturbs p27 translation control and contributes to pituitary tumorigenesis. Cancer research 70: 6026–6035.

Boultwood J, Pellagatti A, Wainscoat JS. 2012. Haploinsufficiency of ribosomal proteins and p53 activation in anemia: Diamond-Blackfan anemia and the 5q- syndrome. Advances in biological regulation 52: 196–203.

Chaligné R, Popova T, Mendoza-Parra MA, Saleem MAM, Gentien D, Ban K, Piolot T, Leroy O, Mariani O, Gronemeyer H et al. 2015. The inactive X chromosome is epigenetically unstable and transcriptionally labile in breast cancer. Genome Research 25: 488–503.

Chen G, Gharib TG, Huang CC, Taylor JM, Misek DE, Kardia SL, Giordano TJ, Iannettoni MD, Orringer MB, Hanash SM et al. 2002. Discordant protein and mRNA expression in lung adenocarcinomas. Molecular & cellular proteomics: MCP 1: 304–313.

De Keersmaecker K. 2015. Ribosomopathies and the paradox of cellular hypo- to hyperproliferation. 125: 1377–1382.

Dreiseitl S, Ohno-Machado L. 2002. Logistic regression and artificial neural network classification models: a methodology review. Journal of biomedical informatics 35: 352–359.

Esposito D, Crescenzi E, Sagar V, Loreni F, Russo A, Russo G. 2014. Human rpL3 plays a crucial role in cell response to nucleolar stress induced by 5-FU and L-OHP. Oncotarget 5: 11737–11751.

Fan H, Li J, Jia Y, Wu J, Yuan L, Li M, Wei J, Xu B. 2017. Silencing of ribosomal protein L34 (RPL34) inhibits the proliferation and invasion of esophageal cancer cells. Oncology research doi:10.3727/096504016x14830466773541.

Fancello L, Kampen KR, Hofman IJ, Verbeeck J, De Keersmaecker K. 2017. The ribosomal protein gene RPL5 is a haploinsufficient tumor suppressor in multiple cancer types. Oncotarget 8: 14462–14478.

Gazda HT, Sheen MR, Vlachos A, Choesmel V, O’Donohue MF, Schneider H, Darras N, Hasman C, Sieff CA, Newburger PE et al. 2008. Ribosomal Protein L5 and L11 Mutations Are Associated with Cleft Palate and Abnormal Thumbs in Diamond-Blackfan Anemia Patients. American Journal of Human Genetics 83: 769–780.

Goudarzi KM, Lindstrom MS. 2016. Role of ribosomal protein mutations in tumor development (Review). International journal of oncology 48: 1313–1324.

Guimaraes JC, Zavolan M. 2016. Patterns of ribosomal protein expression specify normal and malignant human cells. Genome Biology 17.

Hong M, Kim H, Kim I. 2014. Ribosomal protein L19 overexpression activates the unfolded protein response and sensitizes MCF7 breast cancer cells to endoplasmic reticulum stress-induced cell death. Biochemical and biophysical research communications 450: 673–678.

Horos R, Ijspeert H, Pospisilova D, Sendtner R, Andrieu-Soler C, Taskesen E, Nieradka A, Cmejla R, Sendtner M, Touw IP et al. 2012. Ribosomal deficiencies in Diamond-Blackfan anemia impair translation of transcripts essential for differentiation of murine and human erythroblasts. Blood 119: 262–272.

Hu G, Chong RA, Yang Q, Wei Y, Blanco MA, Li F, Reiss M, Au JS, Haffty BG, Kang Y. 2009. MTDH Activation by 8q22 Genomic Gain Promotes Chemoresistance and Metastasis of Poor-Prognosis Breast Cancer. Cancer cell 15: 9–20.

Jung Y, Lee S, Choi HS, Kim SN, Lee E, Shin Y, Seo J, Kim B, Jung Y, Kim WK et al. 2011. Clinical validation of colorectal cancer biomarkers identified from bioinformatics analysis of public expression data. Clinical cancer research: an official journal of the American Association for Cancer Research 17: 700–709.

Kardos GR, Dai MS, Robertson GP. 2014. Growth Inhibitory Effects of Large Subunit Ribosomal Proteins in Melanoma. Pigment cell & melanoma research 27: 801–812.

Khan FH, Pandian V, Ramraj S, Natarajan M, Aravindan S, Herman TS, Aravindan N. 2015. Acquired genetic alterations in tumor cells dictate the development of high-risk neuroblastoma and clinical outcomes. BMC Cancer 15.

Kobayashi T, Sasaki Y, Oshima Y, Yamamoto H, Mita H, Suzuki H, Toyota M, Tokino T, Itoh F, Imai K et al. 2006. Activation of the ribosomal protein L13 gene in human gastrointestinal cancer. International journal of molecular medicine 18: 161–170.

Koussounadis A, Langdon SP, Um IH, Harrison DJ, Smith VA. 2015. Relationship between differentially expressed mRNA and mRNA-protein correlations in a xenograft model system. Scientific reports 5: 10775.

Lai MD, Xu J. 2007. Ribosomal Proteins and Colorectal Cancer. Current genomics 8: 43–49.

Landry DM, Hertz MI, Thompson SR. 2009. RPS25 is essential for translation initiation by the Dicistroviridae and hepatitis C viral IRESs. Genes & Development 23: 2753–2764.

Muhs M, Yamamoto H, Ismer J, Takaku H, Nashimoto M, Uchiumi T, Nakashima N, Mielke T, Hildebrand PW, Nierhaus KH et al. 2011. Structural basis for the binding of IRES RNAs to the head of the ribosomal 40S subunit. Nucleic acids research 39: 5264–5275.

Naora H, Takai I, Adachi M, Naora H. 1998. Altered cellular responses by varying expression of a ribosomal protein gene: sequential coordination of enhancement and suppression of ribosomal protein S3a gene expression induces apoptosis. The Journal of cell biology 141: 741–753.

Noller HF, Hoffarth V, Zimniak L. 1992. Unusual resistance of peptidyl transferase to protein extraction procedures. Science (New York, NY) 256: 1416–1419.

Ormandy CJ, Musgrove EA, Hui R, Daly RJ, Sutherland RL. 2003. Cyclin D1, EMS1 and 11q13 amplification in breast cancer. Breast cancer research and treatment 78: 323–335.

Paquet ER, Hovington H, Brisson H, Lacombe C, Larue H, Tetu B, Lacombe L, Fradet Y, Lebel M. 2015. Low level of the X-linked ribosomal protein S4 in human urothelial carcinomas is associated with a poor prognosis. Biomarkers in medicine 9: 187–197.

Parris TZ, Kovacs A, Hajizadeh S, Nemes S, Semaan M, Levin M, Karlsson P, Helou K. 2014. Frequent MYC coamplification and DNA hypomethylation of multiple genes on 8q in 8p11-p12-amplified breast carcinomas. Oncogenesis 3: e95.

Ruggero D, Shimamura A. 2014. Marrow failure: a window into ribosome biology. Blood 124: 2784–2792.

Russo A, Russo G. 2017. Ribosomal Proteins Control or Bypass p53 during Nucleolar Stress. International Journal of Molecular Sciences 18.

Russo A, Saide A, Cagliani R, Cantile M, Botti G, Russo G. 2016. rpL3 promotes the apoptosis of p53 mutated lung cancer cells by down-regulating CBS and NFκB upon 5-FU treatment. Scientific reports 6.

Russo A, Saide A, Smaldone S, Faraonio R, Russo G. 2017. Role of uL3 in Multidrug Resistance in p53-Mutated Lung Cancer Cells. International Journal of Molecular Sciences 18.

Shenoy N, Kessel R, Bhagat TD, Bhattacharyya S, Yu Y, McMahon C, Verma A. 2012. Alterations in the ribosomal machinery in cancer and hematologic disorders. Journal of Hematology & Oncology 5: 32.

Shi C, Wang Y, Guo Y, Chen Y, Liu N. 2017. Cooperative down-regulation of ribosomal protein L10 and NF-kappaB signaling pathway is responsible for the anti-proliferative effects by DMAPT in pancreatic cancer cells. Oncotarget 8: 35009–35018.

Sim EU, Chan SL, Ng KL, Lee CW, Narayanan K. 2016. Human Ribosomal Proteins RPeL27, RPeL43, and RPeL41 Are Upregulated in Nasopharyngeal Carcinoma Cell Lines. Disease markers 2016: 5179594.

Spatz A, Borg C, Feunteun J. 2004. X-chromosome genetics and human cancer. Nature reviews Cancer 4: 617–629.

Sridhar K, Ross DT, Tibshirani R, Butte AJ, Greenberg PL. 2009. Relationship of differential gene expression profiles in CD34(+) myelodysplastic syndrome marrow cells to disease subtype and progression. Blood 114: 4847–4858.

Sun XX, Dai MS, Lu H. 2007. 5-fluorouracil activation of p53 involves an MDM2-ribosomal protein interaction. The Journal of biological chemistry 282: 8052–8059.

Taghavi A, Akbari ME, Hashemi-Bahremani M, Nafissi N, Khalilnezhad A, Poorhosseini SM, Hashemi-Gorji F, Yassaee VR. 2016. Gene expression profiling of the 8q22-24 position in human breast cancer: TSPYL5, MTDH, ATAD2 and CCNE2 genes are implicated in oncogenesis, while WISP1 and EXT1 genes may predict a risk of metastasis. Oncology Letters 12: 3845–3855.

Tian Q, Stepaniants SB, Mao M, Weng L, Feetham MC, Doyle MJ, Yi EC, Dai H, Thorsson V, Eng J et al. 2004. Integrated genomic and proteomic analyses of gene expression in Mammalian cells. Molecular & cellular proteomics: MCP 3: 960–969.

van der Maaten LJPH, G. E.. 2008. Visualizing High-Dimensional Data Using t-SNE. Journal of Machine Learning Research 9: 2579–2605.

Vogazianou AP, Chan R, Bäcklund LM, Pearson DM, Liu L, Langford CF, Gregory SG, Collins VP, Ichimura K. 2010. Distinct patterns of 1p and 19q alterations identify subtypes of human gliomas that have different prognoses(). Neuro-Oncology 12: 664–678.

Warner JR, McIntosh KB. 2009. How common are extraribosomal functions of ribosomal proteins? Molecular cell 34: 3–11.

Xue S, Barna M. 2012. Specialized ribosomes: a new frontier in gene regulation and organismal biology. Nature reviews Molecular cell biology 13: 355–369.

Yelick PC, Trainor PA. 2015. Ribosomopathies: Global process, tissue specific defects. Rare diseases (Austin, Tex) 3: e1025185.

Yong WH, Shabihkhani M, Telesca D, Yang S, Tso JL, Menjivar JC, Wei B, Lucey GM, Mareninov S, Chen Z et al. 2015. Ribosomal Proteins RPS11 and RPS20, Two Stress-Response Markers of Glioblastoma Stem Cells, Are Novel Predictors of Poor Prognosis in Glioblastoma Patients. PloS one 10: e0141334.

Yuan BZ, Zhou X, Zimonjic DB, Durkin ME, Popescu NC. 2003. Amplification and overexpression of the EMS 1 oncogene, a possible prognostic marker, in human hepatocellular carcinoma. The Journal of molecular diagnostics: JMD 5: 48–53.

Zhou X, Liao WJ, Liao JM, Liao P, Lu H. 2015. Ribosomal proteins: functions beyond the ribosome. Journal of Molecular Cell Biology 7: 92–104.

